# Variant filters using segregation information improve mapping of nectar-production genes in sunflower (*Helianthus annuus* L.)

**DOI:** 10.1101/2024.12.03.626666

**Authors:** Ashley C. Barstow, James P. McNellie, Brian C. Smart, Kyle G. Keepers, Jarrad R. Prasifka, Nolan C. Kane, Brent S. Hulke

**Affiliations:** Department of Plant Sciences, North Dakota State University, Fargo, 58102, ND, United States; Sunflower Improvement Research Unit, Edward T. Schafer Agricultural Research Center, United States Department of Agriculture (USDA)-Agricultural Research Service, Fargo, 58102, ND, United States; Ecology and Evolutionary Biology Department, University of Colorado, Boulder, 80309, CO United States

## Abstract

Accurate variant calling is critical for identifying the genetic basis of complex traits, yet filters used in variant detection and validation may inadvertently exclude valuable genetic information. In this study, we compare common sequencing depth filters, used to eliminate error-prone variants associated with repetitive regions and technical issues, with a biologically relevant filtering approach that targets expected population-level Mendelian segregation. The resulting variant sets were evaluated in the context of nectar volume QTL mapping in sunflower (*Helianthus annuus* L.). Our previous research failed to detect a significant interval containing a strong candidate gene for nectar production (*HaCWINV2)*. We removed certain hard filters and implemented a Chi-square goodness-of-fit test to retain variants that segregate according to expected genetic ratios. We hypothesized that this will enhance mapping resolution and capture key genetic regions previously missed. We demonstrate that biologically relevant filtering retains more significant QTL and candidate genes, including *HaCWINV2*, while removing variants due to technical errors more effectively, and accounted for 48.55% of phenotypic variation. In finding nine putative homologs of *Arabidopsis* genes with nectary function within QTL regions, we demonstrate that this filtering strategy, which considers biological contexts, has a higher power of true variant detection than the commonly used variant depth filtering strategy.

**PLAIN LANGUAGE SUMMARY:** In genomic research, identifying genetic markers is key to understanding complex traits, but traditional methods for filtering genetic data can sometimes miss important information. In this study, we explored a new data filtering approach for mapping genes related to nectar production in sunflower. We applied a more flexible filtering method that considers how markers are expected to segregate in breeding populations. Our previous work failed to identify an important gene previously hypothesized to be involved in nectar production, likely due to overly strict filtering. Our improved approach identified nine sunflower genes related to nectar production genes in the model species *Arabidopsis thaliana*, as compared to zero genes identified from the previous filtering strategy. This study highlights the value of using flexible, biologically relevant filtering methods, which can lead to better results in plant genomic studies.

**CORE IDEAS:**

- Discovering biologically meaningful variants from sequence data requires a careful and critical view of bioinformatic workflows.
- The use of arbitrary filters can remove significant genomic variation that contributes to the phenotype of interest.
- Arbitrary filters can also fail to remove variant call errors.
- A Chi-square filtering strategy based on segregation ratio retained a larger number of valid variants.
- More candidate regions with putative nectar-related genes and better statistical support were discovered.

## 1 INTRODUCTION

Accurate and reliable variant calling is essential for understanding the genetic underpinnings of complex traits. Genomic data possesses errors that present challenges to discovering “true” variants, which often necessitate multiple data filtering strategies to remove. For example, the Genome Analysis Toolkit (GATK) includes a range of tunable parameters to remove low-quality variants and artifacts, which are typically referred to as hard filters (De Summa et al., 2017). These hard filters involve setting specific quality parameter thresholds that variants must meet to be included in downstream analysis. While hard filters help reduce noise and false positives, they can also inadvertently exclude valuable genetic information, potentially affecting the power and resolution of genetic studies. This became apparent in our initial study of quantitative trait loci (QTL) linked to nectar volume in sunflower (Barstow et al., 2022). Although significant QTL were identified, a sunflower homolog (cell wall invertase*, HaCWINV2*) of a well-studied *Arabidopsis thaliana* gene, *CWINV4*, both with evidence of involvement in nectar production, was not detected (Ruhlmann et al., 2010; Prasifka et al., 2018; Minami et al., 2021). We hypothesized that the limited number of variants, due to overly stringent filtering, might have prevented us from capturing key genetic regions. Relaxing hard filters, while retaining variants that segregate in a Mendelian fashion, could lead to improved mapping resolution. Conversely, using erroneous variants in mapping can obscure true genetic relationships and linkage patterns, leading to genetic map inflation, inaccurate mapping of QTL, and misinterpretation of gene interactions (Shields et al, 1991; Buetow, 1991; Lorieux et al., 1995; Lu et al., 2002; Hackett & Broadfoot, 2003).

There is no “one size fits all” approach to selecting parameters and algorithms for modern genomics protocols, as organisms’ molecular genetic characteristics—such as genome size, proportion of repetitive sequences, and presence of structural variants—differ greatly. As our understanding of genomes and technology improves, variant detection methods and quality filters must adapt to the specific attributes of the sequencing data to ensure errors are removed with minimal loss of biological information. Unlike arbitrary thresholds based on depth of coverage, biologically relevant filters that focus on the Mendelian segregation of variants could offer a more meaningful approach to variant quality filtering. We hypothesize that variants that segregate according to expected genetic ratios are more likely to be biologically informative. Furthermore, erroneous variants that deviate from population-level segregation ratios will naturally be excluded through this filtering. The key advantage is that these decisions are based on biological relevance rather than rigid technical thresholds.

Nectar production is a complex trait of interest as hybrid sunflower seed production is fully dependent on pollinators (Greenleaf and Kremen, 2006). Nectar is offered as a reward to increase pollinator visitation, leading to enhanced pollination and increased seed yield in stressful seed and commercial production environments (Simpson and Neff, 1983; Greenleaf and Kremen, 2006; Prasifka et al., 2018). The genetic architecture of nectar production in plants is complex, involving multiple genes that operate within different pathways including carbohydrate metabolism, sugar transport, hormonal regulation, and developmental processes (Reeves et al., 2012; Lee et al., 2005; Seo et al., 2001; Bender et al., 2013; Schmitt et al., 2018). Discovering loci associated with nectar production will allow breeders to more effectively make selections via variant assisted selection and improve yield by increasing pollination rates.

In this paper, we propose an alternative to stringent hard filters previously used in Barstow et al. (2022). Single nucleotide polymorphic (SNP) variants were filtered using the Chisquare goodness of fit test with the expected Mendelian ratio assumed from the filial generation of the population and no known selection. Previously, a Chi-square filter was used in addition to single copy filters, quality filters, minor allele frequency filters, and maximum missingness filters, which results in a limited number of variants (Reinert et al., 2019; Pogoda et al., 2020; Barstow et al., 2022). We compare the results from different filtering approaches on QTL mapping results to illustrate the importance of a more nuanced curation of sequencing data.

## 2 MATERIALS AND METHODS

### 2.1 Experimental Design

The experimental design, including selection of plant materials, controlled-environment phenotyping, in-field validation of phenotypes and genotyping methods and materials are reported in Barstow et al. (2022). Briefly, the parental lines used to create the mapping population had contrasting nectar volume and sucrose content (Mallinger and Prasifka, 2017). HA 434 has high oleic (HO) acid in the seed oil, and a relatively high volume of dilute, hexose-rich (glucose and fructose) nectar while HA 456 has HO seed oil and a lower volume of nectar with an unusually high concentration of sucrose (Miller et al., 2004; Miller et al., 2006). HA 456 is an inbred line derived from HA 434 crossed with S-16 YU (Miller et al., 2006). The high selection pressure for the HO acid haplotype in the development of parent lines HA 434 and HA 456 resulted in HA 456 inheriting the haplotype associated with *FAD2-1* from HA 434 (Schuppert et al., 2006). The mapping population was founded from a single F_1_ individual and underwent single seed descent until F_6_ seeds of 198 recombinant inbred lines were produced.

### 2.2 Data Curation

Demultiplexed data was downloaded directly from Novogene’s servers (Novogene, Sacramento, CA). From the demultiplexed data, three datasets were developed in this experiment. The first dataset was made with only hard filters and no additional Chi-square filtering. The second dataset was the variant set from the initial nectar volume QTL mapping study (Barstow et al., 2022) with both hard filters and Chi-square filtering as described below. The third was the experimental dataset curated without hard filters and only the population-level, segregation ratio filter, based on a Chi-square goodness of fit test.

Our first dataset was developed using the following hard filters. Raw genomic libraries were trimmed using Trimmomatic Version 0.38 (Bolger et al., 2014) using the code: NexteraPE-PE.fa:2:30:10 LEADING:3 TRAILING:3 MINLEN:100, with NexteraPE-PE.fa containing the standard set of Nextera adapters to be trimmed from reads. The resulting trimmed reads were aligned to a *Helianthus annuus* chromosome-scale reference genome, HA 412 HOv2.0 (Badouin et al., 2017). Variant calling was performed using GATK best practices (Van der Auwera et al., 2013; Van der Auwera & O’Connor, 2020), resulting in a single variant call file (vcf). This vcf table was then filtered for single copy sites based on depth (sites with depth across all sunflower lines [n = 192] between 200-1000 were retained). This range of depths was selected by creating a histogram of the variant depths in the vcf table, which produces a prominent, normally distributed peak at the mode single-copy depth sampling of the summed aligned libraries. Variants at depths that are too low were discarded for possibly representing sequencing errors, whereas variants at excessively high depths were discarded for potentially belonging to repetitive content, which is a major contributor of noise to a dataset. Additional filtering was done to select variants above a minimum quality score of 100 (minQ=100), minor allele frequency of 0.05 or greater (maf=0.05) and no more than 25% of samples having missing data for a given variant (maxmissing=0.75). Missing data were imputed using BEAGLE version 5.0 with default settings retained (Browning et al., 2018).

Our second approach was the original variant set from a previous nectar volume QTL mapping study (Barstow et al., 2022). All previous steps from the first set were followed. Following imputation, polymorphic SNPs were then filtered using a custom script invoking PROC FREQ of SAS v. 9.4 (SAS Institute, 2016) to exclude variants that did not fit the expected F_6_ segregation ratio, from a Chi-square analysis goodness of fit test (p > 0.10). This dataset was curated using both a Chi-square filter and hard filters.

The third approach using Chi-square filtering was prepared as follows. First, *fastp* was used to perform quality control and exclude adapters (Chen et al., 2018). Using BWA-mem2, FASTQ files were aligned with the most recent sunflower genome assembly, HA 412 HO v2.0 (Vasimuddin et al., 2019; Badouin et al., 2017). None of the previously described filters (minQ, maf, max-missing, or variant-depth) were used. Missing data were imputed using BEAGLE version 5.4 with default settings retained (Browning et al., 2018). Polymorphic SNPs were then filtered using a custom R script to exclude variants that did not fit the expected F_6_ segregation ratio, from a Chi-square goodness of fit test (p > 0.10). The custom R script produces identical output to the PROC FREQ script used previously but had improved run time with larger variant sets.

### 2.3 Linkage Mapping and QTL Analysis

For the hard filtered dataset and the Chi-square filtered dataset (first and third approaches), a custom R script was used to prune additional SNPs based on distance, aiming to reduce variant numbers to meet software requirements for QTL mapping. For each chromosome, the distance between sets of three variants was calculated and if the distance was less than 125,000 base pairs, one is randomly selected to be kept; otherwise, it keeps all variants in that range (*marker_filt_dist*; Supplemental Material). Additionally, for all datasets, a custom R script was used to remove redundant variants by comparing variants at the same genetic map position and retaining the SNP with the higher p-value from Chi-square test of Mendelian segregation (*thinning_loop*; Supplemental Material).

The construction of linkage maps for all datasets were carried out in R/qtl package version 1.60 (Broman et al., 2003). Genetic distances were calculated using the Kosambi map function (Kosambi, 1944). Erroneous SNPs were identified and manually removed based on recombination maps and SNP frequencies. For the QTL analysis, the function *scanone* was employed to identify putative QTL with datasets that created a biologically feasible linkage map. Additionally, two-dimensional scans were conducted using *scantwo* with Haley-Knott regression (Haley and Knott, 1992) to explore interactions between QTL. The thresholds were established based on the results of 1,000 permutations at a p=0.05 significance level (Broman et al., 2003).

Building upon the results from single QTL model and the two-dimensional genome scan, multiple QTL models (MQM) were fit to examine the presence of additional QTL and QTL-by-QTL interactions as described by Broman and Sen (2009). QTL above the permutation threshold formed the initial model and additional model terms were discovered using the *addqtl* and *addint* functions. The resulting final MQM model incorporated all significant QTL and interactions above a LOD score of 3. The effects of individual QTL were evaluated by comparing the full model to one where the respective term was omitted. LOD scores, estimated additive effects, and the percentage of phenotypic variance explained by each QTL and QTL interaction were obtained from the drop-one analysis of variance (ANOVA) table. Using the resulting model, an additional ANOVA was used to verify results.

### 2.4 Candidate gene analysis

Using previous knowledge of the cloned genes in *Arabidopsis* with implicated functions in nectaries and nectars (Table 2 from Roy et al., 2017), the corresponding protein sequences were queried against the reference genome HA 412 HO v2.0 (Badouin et al. 2017) using tblastn as implemented in BLAST+ 2.11.0 (Altschul et al., 1990). The protein sequences were obtained from The Arabidopsis Information Resource (TAIR) website (https://www.arabidopsis.org/). The candidate genes were considered if found within the 2.0 LOD drop interval of the identified loci. The corresponding candidate gene nucleotide sequences were extracted, and the functional annotation was cross-checked with a TBLASTX 2.16.0+ query against the core_nt database. Those genes that retained putative functional characterization, as determined by best result, were retained. An additional search was conducted with the most up-to-date annotation of HA 412 HO v2.0 (Kyle Keepers, unpublished data) to determine the total number of genes within each QTL region.

## 3 RESULTS

### 3.1 Construction of Linkage Maps

The hard filtered dataset had 289,678 variants, which is beyond the capacity for most linkage mapping software to efficiently analyze. For mapping efficiency, redundant and erroneous variants were removed, as previously described, resulting in 7425 variants. The resulting linkage map had substantially inflated genetic distances that exceed biologically plausible lengths. The number of variants on each chromosome ranged from 362 to 538 with genetic distances ranging from 43808.3 to 65346.4 centimorgans (cM). Due to the relationship between the parental lines used to create the mapping population, we expected that there would be genomic regions with conserved haplotypes between the parents and no variant segregation, leading to clustering of SNPs across the genome. However, this was not observed as the linkage map had limited clustering; SNPs were dispersed across all 17 chromosomes, including chromosome 14 despite both parents having the same HO haplotype associated with the *FAD2-1* locus from high selection pressure (Miller et al., 2006; Schuppert et al., 2006).

For the dataset with both filters (Chi-square and hard filters), the variant set started with 765 variants spanning 16 different chromosomes, excluding chromosome 14 due to a lack of polymorphic SNPs-(for reasons already described). Chromosome 6 only included one variant, which was excluded from the linkage map. The remaining number of variants ranged from two to 124 per chromosome, forming localized clusters on each due to large segments of the genomes of the two parents being identical by descent. Additional erroneous variants were identified based on the recombination map and manually dropped, resulting in a linkage map containing 748 variants over 15 chromosomes. Gaps of 27.9, 25.9, 38.7, 34.0, and 26.8 cM were observed on chromosomes 3, 5, 11, 12, and 15, respectively. The average variant density by chromosome ranged from 0.3 to 19.7 variants per cM due to the low-density nature of the map.

The Chi-square filtered dataset had 1,151,856 SNPs. Similarly, for mapping efficiency, redundant and erroneous variants were removed, resulting in 3631 variants. The variants span across 16 chromosomes, excluding chromosome 14. The number of variants per chromosome ranged from 42 to 479, and the average variant density by chromosome ranged from 0.5 to 5.2 variants per cM with an overall average of 1.4. Chromosome length ranged from 159.5 cM to 667.1 cM.

### 3.2 QTL analysis of nectar volume

QTL analysis was not performed on the hard filtered dataset because of the poor quality of the linkage map.

Using the dataset with both filters (Chi-square and hard filters), no significant loci were identified using the single-QTL model. However, the variant with the highest LOD score (2.59), although not statistically significant, mapped to chromosome 16 and was within the region identified as the largest main effect QTL in Barstow et al., 2022. With no significant QTL, MQM was not performed for the dataset using hard filters.

For the Chi-square filtered dataset, the QTL analysis identified nine significant QTL and three QTL interactions contributing to the trait variation in the dataset. The full model, developed using Haley-Knott regression and model fitting, explained 48.55 % of the phenotypic variance (LOD = 27.42, p-value < 0.001). When dropping one QTL at a time, the most significant QTL was identified on chromosome 10 at 402.9 cM (LOD =12.64, 18.44 % variance explained; Table 1). Overall, the main effect QTL identified ranged from LOD of 3.17 to 12.64. Three QTL interactions were detected: chromosome 10 at 402.9 cM with chromosome 11 at 3.2 cM (LOD = 10.47, 14.86 % variance explained), chromosome 10 at 402.9 cM with chromosome 16 at 32.9 cM (LOD = 3.40, 4.42 % variance explained), and chromosome 15 at 16.2 cM with chromosome 16 at 32.9 cM (LOD = 7.07, 9.61 % variance explained).

**Table 1.**
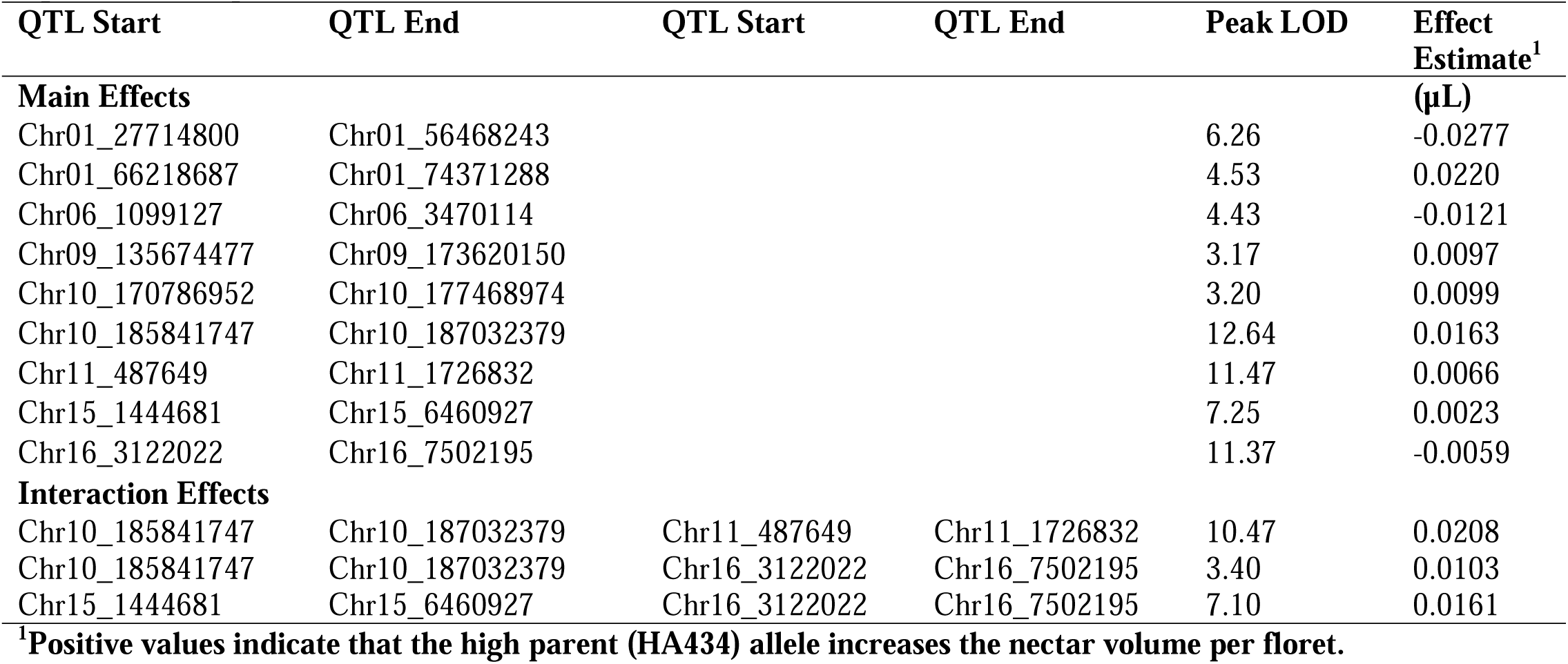
Summary of QTL effects on the nectar volume trait in sunflower from dataset with only Chisquare filtering.

| QTL Start | QTL End | QTL Start | QTL End | Peak LOD | Effect Estimate<sup>1</sup> |
| --- | --- | --- | --- | --- | --- |
| **Main Effects** | | | | | ($\mu\text{L}$) |
| Chr01_27714800 | Chr01_56468243 |  |  | 6.26 | -0.0277 |
| Chr01_66218687 | Chr01_74371288 |  |  | 4.53 | 0.0220 |
| Chr06_1099127 | Chr06_3470114 |  |  | 4.43 | -0.0121 |
| Chr09_135674477 | Chr09_173620150 |  |  | 3.17 | 0.0097 |
| Chr10_170786952 | Chr10_177468974 |  |  | 3.20 | 0.0099 |
| Chr10_185841747 | Chr10_187032379 |  |  | 12.64 | 0.0163 |
| Chr11_487649 | Chr11_1726832 |  |  | 11.47 | 0.0066 |
| Chr15_1444681 | Chr15_6460927 |  |  | 7.25 | 0.0023 |
| Chr16_3122022 | Chr16_7502195 |  |  | 11.37 | -0.0059 |
| **Interaction Effects** |  |  |  |  |  |
| Chr10_185841747 | Chr10_187032379 | Chr11_487649 | Chr11_1726832 | 10.47 | 0.0208 |
| Chr10_185841747 | Chr10_187032379 | Chr16_3122022 | Chr16_7502195 | 3.40 | 0.0103 |
| Chr15_1444681 | Chr15_6460927 | Chr16_3122022 | Chr16_7502195 | 7.10 | 0.0161 |

<sup>1</sup>Positive values indicate that the high parent (HA434) allele increases the nectar volume per floret.

**Table 2.** Homologous nectar-production related candidate genes of *Arabidopsis thaliana* genes within sunflower genome.

| Candidate Gene (*Arabidopsis thaliana*) | Function and/or mutant phenotype | Sunflower QTL |  |  | E-value | Bitscore |
| --- | --- | --- | --- | --- | --- | --- |
|  |  | Chromosome | Start Location | End Location |  |  |
| *ARF6* | Transcription factor; mutants lack nectaries (Reeves et al., 2012) | Chr09 | 146,751,503 | 146,751,357 | 2.22E-05 | 54.7 |
|  |  | Chr11 | 1,078,087 | 1,077,932 | 1.12E-16 | 92 |
| *ARF8* | Transcription factor; mutants lack nectaries (Reeves et al., 2012) | Chr09 | 149,615,897 | 149,615,769 | 9.00E-08 | 55.8 |
|  |  | Chr11 | 1,078,087 | 1,077,932 | 8.00E-17 | 92.4 |
| *CRABS CLAW* | Transcription factor needed for nectary development (Bowman & Smyth, 1999) | Chr06 | 1,332,589 | 1,332,362 | 2.45E-08 | 58.5 |
| *CWINV4* | Cell wall invertase responsible for nectar production; mutants do not produce nectar (Ruhmann et al., 2010) | Chr01 | 68,660,146 | 68,659,286 | 5.55E-109 | 365 |
| *MYB21* | Transcription factor; mutants lack nectar and expression of SWEET9 (Schmitt et al., 2018) | Chr09 | 153,815,731 | 153,815,432 | 4.70E-22 | 99.8 |
|  |  | Chr09 | 146,202,208 | 146,201,897 | 3.27E-21 | 96.7 |
| *MYB57* | Transcription factor; mutants have smaller nectaries and produce about half the nectar of wild type (Bender et al., 2013) | Chr06 | 1,517,950 | 1,518,204 | 1.48E-25 | 109 |
|  |  | Chr10 | 175,680,709 | 175,680,380 | 1.98E-21 | 96.7 |
| *PIN6* | Transmembrane protein involved in auxin transport; mutants produced 60% less nectar (Bender et al., 2013) | Chr01 | 54,495,305 | 54,495,153 | 2.66E-17 | 63.5 |
|  |  | Chr09 | 136,142,804 | 136,142,373 | 1.07E-32 | 140 |
| *SWEET9* | Sucrose uniporter; mutants do not produce nectar (Lin et al., 2014) | Chr09 | 170,112,792 | 170,113,016 | 5.84E-04 | 46.2 |
|  |  | Chr15 | 4,825,680 | 4,825,889 | 1.04E-14 | 79.3 |
| *TIR1* | Auxin receptor that mediates auxin-regulated transcription (Schmitt et al., 2018) | Chr15 | 4,842,038 | 4,841,253 | 3.35e-127 | 418 |

The phenotypic range of nectar volume in the mapping population florets spans 0.03 µl/floret to 0.91 µl/floret in the 192 entries sampled, exhibiting similar nectar volumes to the parents, but also with values that appeared intermediate (Figure 1 from Barstow et al., 2022). The estimated additive effects of individual QTL ranged from −0.0277 µl/floret (chromosome 1 at 88.0 cM) to 0.0220 µl/floret (chromosome 1 at 191.0 cM). Positive effect estimates indicate that the allele contributed by the high parent (HA 434) increased nectar volume per floret and negative effects meaning the low parent (HA456) contributed the allele increasing nectar volume. The interaction effects ranged from 0.0103 to 0.0208 µl/floret. QTL identified in this study, though modest in individual contributions, holds biological relevance when considered collectively and within the broader phenotypic range.

**Figure 1:**
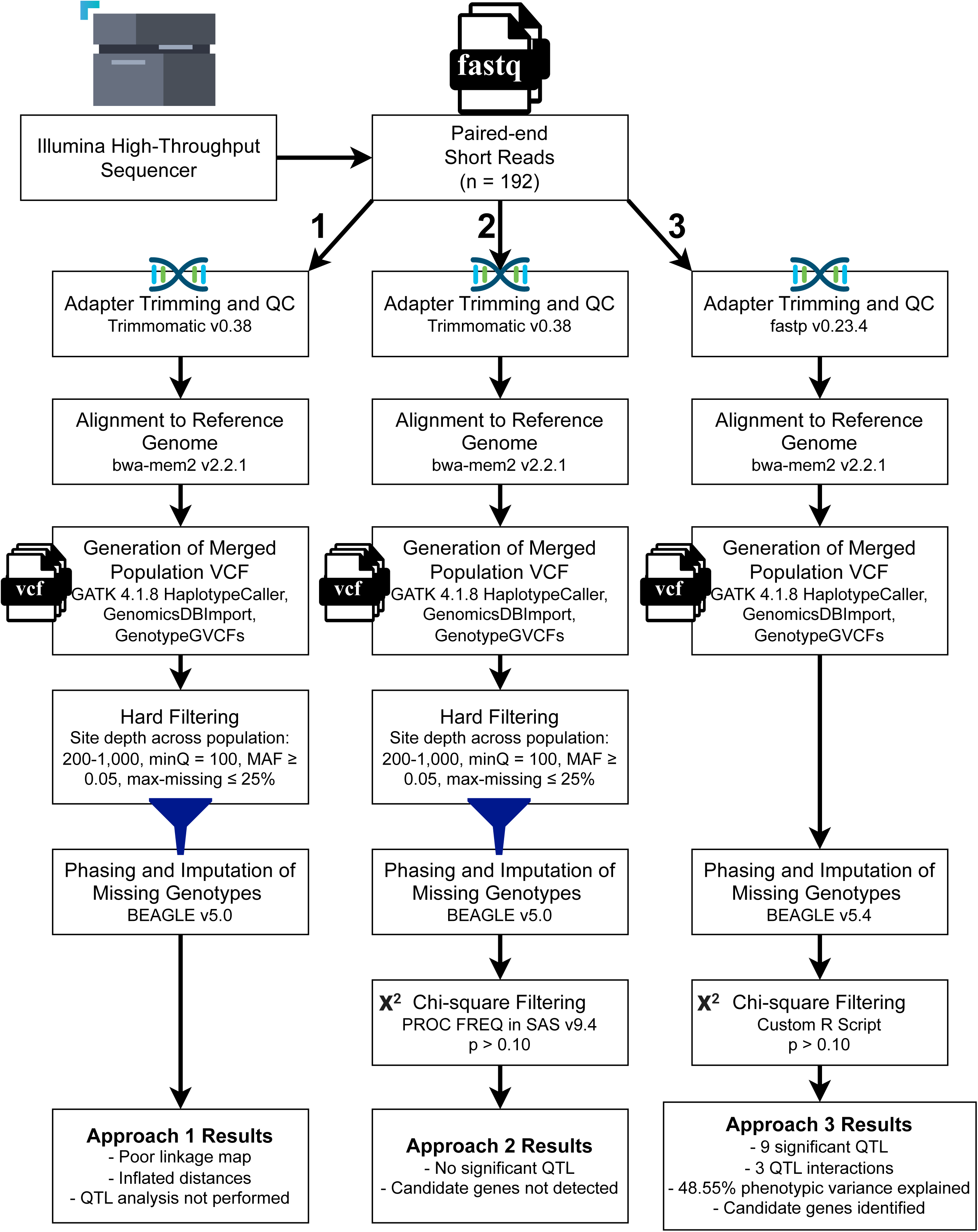
Procedural comparison of three VCF filtering approaches for sunflower genomic data analysis, with a synopsis of results.

### 3.3 Candidate Genes

Of the nectar-related genes listed in Roy et al. (2017), 9 homologous genes were found within the regions identified from this study with blast bit scores ranging from 46.2 to 418: *CWINV4, SWEET9, PIN6, MYB57, MYB21, CRABS CLAW, ARF6, ARF8, TIR1* (Table 2). Based on the blast results, the most similar sunflower gene to an *Arabidopsis* homolog was a *TIR1* homolog found within the QTL region on chromosome 15 (e-value = 3.35e-127). Notably, a *CWINV* was found within the QTL region on chromosome 1 (e-value = 5.55e-109) as previously hypothesized from Prasifka et al. (2018). Only candidate genes whose functional annotations matched the corresponding *Arabidopsis* genes were reported (Table 2).

An additional search was conducted with the most up-to-date annotation of HA 412 HOv2.0 to determine the number of genes within each QTL region (within 2 LOD units). The number of annotated genes within each QTL interval ranged from 29 to 839 genes (Table 3).

**Table 3.** Additional candidate genes discovered within QTL 2 LOD unit interval on the HA412HO v2.0 sunflower genome.

| **Chromosome** | **Start Location** | **End Location** | **Length (bp)** | **No. of Genes** |
| --- | --- | --- | --- | --- |
| Chr01 | 27,714,800 | 56,468,243 | 28,753,443 | 277 |
| Chr01 | 66,218,687 | 74,371,288 | 8,152,601 | 102 |
| Chr06 | 1,099,127 | 3,470,114 | 2,370,987 | 59 |
| Chr09 | 135,674,477 | 173,620,150 | 37,945,673 | 839 |
| Chr10 | 170,786,952 | 177,468,974 | 6,682,022 | 165 |
| Chr10 | 185,841,747 | 187,032,379 | 1,190,632 | 29 |
| Chr11 | 487,649 | 1,726,832 | 1,239,183 | 49 |
| Chr15 | 1,444,681 | 6,460,927 | 5,016,246 | 202 |
| Chr16 | 3,122,022 | 7,502,195 | 4,380,173 | 121 |

## 4 DISCUSSION

The challenges associated with variant calling and filtering highlight the need for more flexible and adaptive approaches in genomic research. We have shown that traditional hard filters inadvertently exclude significant genetic variation that contributes to phenotypic traits. By utilizing goodness-of-fit tests based on Mendelian ratios, such as the Chi-square test, rather than relying on stringent hard filters, we were able to ensure a more accurate and comprehensive representation of the genetic architecture, preserving valuable variants that may be crucial for QTL mapping and downstream analyses. Conversely, the Chi-square filter helped ensure unreliable variants were filtered out, which did not occur completely in our dataset with standard hard filters. This shift in filtering strategy offers the potential to enhance the resolution and power of genetic studies, particularly in complex genomes like that of sunflower and polygenic traits such as nectar production.

Using arbitrary thresholds in hard filters removed significant genomic variation that contributed to the trait of interest: nectar volume in sunflower. The use of MAF, quality scores, and a local sequencing depth filter in addition to a Chi-square filter of the segregation ratio of that population, retained a limited number of genetic variants, 746 SNPs over 15 chromosomes. While in Barstow et al., 2022, significant QTLs were identified using that variant set and a different mapping algorithm, using the mapping algorithms and methods described herein, we were unable to identify significant regions. However, after removing hard filters and considering only the Mendelian segregation patterns of that generation to filter the variants, we generated over 1 million SNPs from which we selected 3631 nonredundant, valid variants spanning across 16 chromosomes for mapping. This approach facilitated the retention of a greater number of variants across the genome, effectively increasing representation in regions that were not covered previously (Figure 2). The improved genome coverage increases the likelihood of capturing loci linked to the phenotype. With those improvements in the genetic map, we were able to capture more candidate regions associated with nectar production with better statistical support than our previous study. The amount of phenotypic variation explained by the model using the Chi-square filtered dataset was 48.55%. The R^2^ value achieved in this study is promising, particularly for a highly quantitative trait like nectar production, as it suggests that a significant portion of the phenotypic variance can be explained by the identified genetic variants despite the complex, polygenic nature of the trait. The overall fit of the model used was highly statistically significant (p < 0.001) and able to identify nine QTL regions and three epistatic interactions while the hard filtered dataset failed to identify any loci with this mapping algorithm above the permutation thresholds. This approach not only improved the genetic map’s resolution but also provided a strong foundation for identifying candidate genes associated with nectar production. Additional variants could have been included post-hoc to increase resolution further in regions of interest.

**Figure 2:**
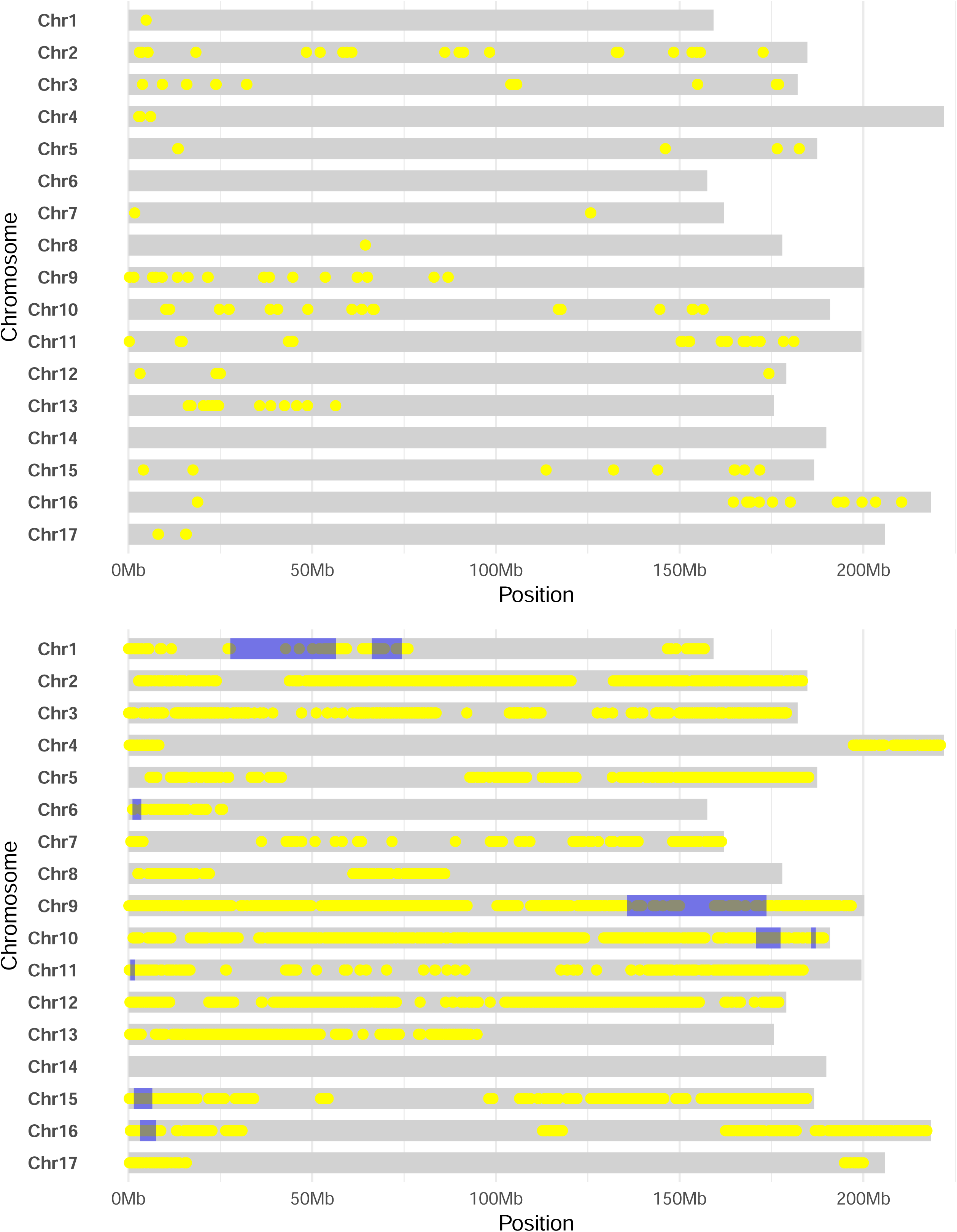
Overview of the physical map of the mapping population, aligned to the HA 412 HOv2 genome. Yellow regions represent polymorphic regions in the genetic map, while blue regions represent QTL regions within 2 LOD units of peak significant variant or loci. (Top) Hard filtered and Chi-square filtered dataset. (Bottom) Chi-square filtered dataset.

Building on the increased genome coverage and the enhanced statistical power of our refined genetic map, we were able to move forward with candidate gene discovery, focusing on pinpointing specific genes within the identified QTL regions that may be involved in nectar production. Building on the previous work done in the model species *Arabidopsis*, candidate gene homologs of *CWINV4, SWEET9, PIN6, MYB21, MYB57, CRABS CLAW, ARF6, ARF8,* and *TIR1* were located within QTL regions (Table 2). Previous studies show that the genetic control of nectar secretion in *Arabidopsis* and *Brassica rapa* is complex, but a key player is cell wall invertase, the enzyme known to catalyze the hydrolysis of sucrose into glucose and fructose (Ruhlmann et al., 2010; Minami et al., 2021). This study was able to identify *HaCWINV2* (the closest homolog to *CWINV4* in the sunflower genome) as a contributor to nectar production, on chromosome 1. Our results support that given the additive and interactive QTL, nectar production in *H. annuus* is polygenic, with some strong candidate genes beneath QTL with large effects (Table 1). The proposed model in Roy et al., 2017 of floral nectary regulation suggests that this process relies on a specific cascade of expression events which ultimately lead to nectar production. For all the genes involved in this proposed pathway (Figure 3 in Roy et al., 2017), we were able to find homologs in this study within 2 LOD units of our identified QTLs, with several regions containing two homologs and two regions containing none from the Roy et al. model. Based on the number of genes within each QTL, strong candidate genes may not be limited to these (Table 3; Supplemental Material). Within the loci with no candidate genes from Roy et al., there were candidate genes with support from other studies. Specifically, within the chromosome 10 (185,841,747 bp 187,032,379 bp) confidence interval: xyloglucan endotransglucosylase/hydrolase, heat shock protein 70, leucine-rich repeat (LRR) protein kinase family protein, glycosyl hydrolases family proteins, zinc finger protein, and Golgi nucleotide sugar transporter (Bowman & Smyth, 1999; Ballerini et al., 2019; Silva et al., 2020). Notable candidate genes found within chromosome 16 (3,122,022 bp −7,502,195 bp) confidence interval included glycosyl hydrolases family proteins, LRR receptor-like serine/threonine-kinase, H[+]-ATPase, zinc finger proteins, transmembrane proteins, and plasma membrane fusion protein (Bowman & Smyth, 1999; Ballerini et al., 2019; Silva et al., 2020).

The utility of this method is currently limited to biparental recombinant inbred line datasets with no selection assumed, in order to properly estimate expected allele frequencies. However, to extend the method to plant breeding datasets and diversity panels, we need to adapt this or similar methods to populations that do not fit these basic assumptions. This work is currently underway in the context of breeding populations with a history of artificial selection for two heterotic groups in hybrid oilseed sunflower.

## 5 CONCLUSION

In this study, three genomic data filtering strategies were compared using nectar-production QTL results in sunflower. Our study revealed that using a Chi-square goodness-of-fit test based on the expected population-level segregation ratio outperformed a standard data curation strategy utilizing hard filters in both retaining valid variants and removing errant variants. Using the Chisquare filtered data without hard filters, a QTL model was developed that was able to explain over 48% of the phenotypic variation while the hard filtered data lacked the ability to identify QTL over the permutation threshold. Nine significant putative QTL and three QTL interactions contributing to the trait variation were found. Additionally, within the identified QTL regions, many genes homologous to previously identified nectar-related genes in other species were found including homologs of *CWINV4*, *SWEET9*, *PIN6*, *MYB21*, *MYB57*, *CRABS CLAW*, *ARF6*, *ARF8*, and *TIR1*. Our results provided us with evidence to carefully reconsider implementation of hard filters in sequencing data curation for genomic studies.

## ABBREVIATIONS

ANOVA: Analysis of variance
BLAST: Basic Local Alignment Search Tool
cM: centimorgans
CWINV: cell wall invertase
GATK: Genome Analysis Toolkit
HA: *Helianthus annuus*
HO: high oleic
IBD: identity-by-descent
LOD: logarithm of the odds
minQ: minimum quality
maf: minor allele frequency
MQM: multiple QTL model
QTL: quantitative trait loci
SNP: single nucleotide polymorphism
Vcf: variant call file format

## SUPPLEMENTAL MATERIAL STATEMENT

One table with the putative functions of all genes known in the QTL LOD support intervals, based on the HA 412 HOv.2 reference genome.

## DATA AVAILABILITY STATEMENT

Data and code described in the methods will be made available in the National Agricultural Library – Ag Data Commons after acceptance, and the DOI provided here.

## AUTHOR CONTRIBUTIONS

Conceptualization, BSH and ACB; Data curation, ACB, BCS, JPM, KGK; Formal analysis, ACB, JPM, JRP; Funding acquisition, BSH, NCK, JRP; Investigation, all authors; Methodology, BCS, ACB, JPM, KGK; Project administration, BSH; Visualization, BCS and ACB; Writing – original draft, ACB; Writing – review & editing, all authors.

## ACKNOWLEDGMENTS

The authors wish to thank Brady Koehler for assisting with plant tissue capture, Dr. Ziv Attia for assisting with sequencing efforts, and numerous undergraduate students that assisted with development of the mapping population.

